# Cryo-EM Structure of a Human LECT2 Amyloid Fibril Reveals a Network of Polar Ladders at its Core

**DOI:** 10.1101/2023.02.08.527771

**Authors:** Logan S. Richards, Maria D. Flores, Samantha Zink, Natalie A. Schibrowsky, Michael R. Sawaya, Jose A. Rodriguez

**Author notes:** Correspondence to Jose A. Rodriguez.

## Abstract

ALECT2 is a type of systemic amyloidosis caused by deposition of the leukocyte cell-derived chemotaxin-2 (LECT2) protein in the form of fibrils. In ALECT2, LECT2 fibril deposits can be found in the glomerulus, resulting in renal failure. Affected patients lack effective treatment options outside of renal transplant or dialysis. While the structure of LECT2 in its globular form has been determined by X-ray crystallography, structures of LECT2 amyloid fibrils remain unknown. Using single particle cryo-EM, we now find that human LECT2 forms robust twisting fibrils with canonical amyloid features. At their core, LECT2 fibrils contain two mating protofilaments, the ordered core of each protofilament spans residues 55-75 of the LECT2 sequence. The overall geometry of the LECT2 fibril displays features in line with other pathogenic amyloids. Its core is tightly packed and stabilized by a network of hydrophobic contacts and hydrogen-bonded uncharged polar residues, while its outer surface displays several charged residues. The robustness of LECT2 fibril cores is illustrated by their limited dissolution in 3M urea and their persistence after treatment with proteinase K. As such, the LECT2 fibril structure presents a potential new target for treatments against ALECT2.

## Introduction

Amyloid diseases are linked to the formation and persistence of large, multimeric structures in various tissues. Amyloid fibrils are characterized by a cross-beta scaffold in which identical protein molecules mate tightly as beta-strands to form a long, unbranched fibril^1,2^. Before amyloid-forming proteins assemble into fibrils, they can display a globular fold, requiring partial or total unfolding to convert into an amyloid state^3,4^. Debilitating neurodegenerative diseases such as Alzheimer’s and Parkinson’s disease are most well-known to involve amyloidogenesis^5,6^, while many other examples of amyloid diseases involve the systemic deposition of persistent amyloid fibrils throughout various organs including the heart, liver, kidneys, skin, digestive tract, and nervous system^7^. While the persistence of amyloid aggregates is a unifying feature of these diseases, the factors contributing to the misfolding, retention and toxicity of each disease-associated fibril are distinct and likely dependent on their structures^8^. This underscores drive to characterize amyloid fibril structures.

In 2008, Benson *et al*. identified a new form of systemic amyloid disease associated with impaired renal function, renal failure and nephrotic syndrome^9^. Biochemical characterization of the fibrils observed in the glomeruli of the kidneys identified the protein leukocyte cell-derived chemotaxin-2 (LECT2) as a major component of the fibrils, inspiring the disease name, ALECT2^9^. Since its identification, ALECT2 has been characterized as a common and likely underdiagnosed form of renal amyloidosis^10^. ALECT2 has been found in high prevalence among patients of Mexican descent in the southwest United States, those of Punjabi descent, First Nations peoples in British Columbia, Egyptians, Chinese of Han ethnicity, and Native Americans^11^. There remains a need for diagnostic tools to distinguish ALECT2 from other renal amyloid diseases in patients and avoid misdiagnosis and improper treatment^12^. In contrast to some other systemic amyloid diseases, there are no available treatments for ALECT2 and no molecules that effectively target the amyloid state of its namesake, LECT2.

Nascent LECT2 is a 151 amino acid polypeptide with an 18-residue N-terminal signal peptide that is cleaved before its secretion into the blood as a globular protein^13^. In circulation, LECT2 performs multiple biological functions. It can act as a chemotactic factor to promote the migration of neutrophils to sites of infection^13,14^. It can also act as a regulator of cartilage growth^15^, and it can act as a hepatokine important for metabolic homeostasis^16^. Structurally, LECT2 displays an M23 metalloendopeptidase fold that coordinates Zn(II), but lacks a critical histidine residue that would enable its catalysis^17^. While the clinical details of ALECT2 disease have been well characterized, the molecular details of LECT2 amyloid aggregation remain unclear. Loss of this bound Zn(II) may play an important role in the conversion of properly folded LECT2 into an amyloid form *in vitro* as it is thought to destabilize a central beta-barrel motif within the protein^18^. Many patients who suffer from ALECT2 are also homozygous for the common I40V sequence polymorph of the protein^19^. This mutation appears to be important, but not sufficient for inducing amyloid formation, and does not destabilize the metal binding properties of the functional protein; accordingly, its role in amyloid conversion remains unclear^18^. Amyloidogenic segments of LECT2 have been identified, and predicted amyloid-forming LECT2 peptides assemble into amyloid-like fibrils in isolation^20^. However, the identity of the LECT2 amyloid core remains unknown.

Recent advances in single-particle cryo-EM using helical reconstruction have permitted the determination of near-atomic resolution structures of a wide variety of amyloid fibrils^3^. These approaches have now allowed us to determine the structure of amyloid fibrils formed by recombinant full-length LECT2. This ~2.7Å resolution cryo-EM structure of a LECT2 fibril reveals a tightly mated amyloid core that spans residues 55 to 75 and harbors a network of polar ladders. Its structure is also a first step toward a more comprehensive molecular understanding of ALECT2 and a starting point for the design of targeted therapeutics or diagnostic agents.

## Results

### Assembly of recombinant human LECT2 amyloid fibrils

Amyloid fibrils were grown from recombinantly expressed, full-length LECT2 protein encoding the I40V sequence polymorph. Purified LECT2 was allowed to assemble into fibrils over a 48-hour period, while shaking at 2400 rpm at a temperature of 37°C. After an initial lag phase, fibrils grew at an exponential rate and then stabilized, as monitored by changes in Thioflavin T (ThT) fluorescence over a 48-hour time course (Figure S1D). Unbranched fibrils were observed by negative stain transmission electron microscopy in samples prepared in an equivalent manner, but without ThT (Figure S1E). Unbranched LECT2 fibrils exhibited a regular helical twist with an ~850Å spacing between crossovers. These features were consistent with other amyloid fibrils, and suggested that LECT2 fibrils harbored a canonical amyloid core supported by steric zipper motifs. Indeed, certain regions of the LECT2 sequence were predicted to have a high propensity to form steric zippers (Figure S1A), and the amyloid nature of the fibrils was confirmed by the appearance of a signature cross-beta diffraction pattern when X-ray diffraction was collected from aligned LECT2 fibrils (Figure S2A).

### LECT2 amyloid fibrils are urea and protease-resistant

Amyloid aggregates are often highly stable and can resist chemical or proteolytic denaturation. To evaluate the stability of LECT2 fibrils, a nephelometric assay was performed that monitored light scattering induced by fibrils in solution. Fibril dissolution was correlated with decreased nephelometry signal and assessed under conditions where fibrils were exposed to either water, fibrillization buffer alone, fibrillization buffer with 3M urea, or 40 nM Proteinase K in 150 mM NaCl, 50 mM MOPS pH 6.5 buffer (Figure S4A). LECT2 fibrils persisted in solution when exposed to fibrillization buffer or water over a 24-hour period. In contrast, exposure to 3M urea induced an immediate, slight decrease in fibril content followed by a stable lower signal over the 24-hour period. Incubation with Proteinase K induced a larger drop in signal over the first two hours of incubation following a stable, lower signal phase during the remaining incubation period. These initial drops in signal could have resulted from the dissolution of larger fibril clumps or the removal of more susceptible fibrils, but in all cases, there remained a stable subset of fibrils in solution (Figure S4B-E). The presence of a species that was resistant to protease digestion correlated with the appearance of a ~7kDa band as observed by SDS PAGE (Figure S9A). The identity of this fragment was investigated 60min after exposing fibrils to the protease using bottom-up mass spectrometry. That analysis identified segments covering 25% of the LECT2 sequence (Supplementary Table 2), including a fragment of the LECT2 fibril core spanning residues N67-R74 and portions of its lysine-rich C-terminal region (Figure S9A).

### Structural determination of a LECT2 amyloid fibril core

After biochemical characterization, LECT2 fibrils were prepared for high-resolution structure determination by single particle cryo-EM. The conditions for grid preparation were optimized to retain regularly twisting fibrils and discourage their adsorption to the air-water interface (Figure S4A and S5). Cryo-electron micrographs of these fibrils produced 2D class averages with fibril characteristics consistent with negative stain images (Figure S3A). Frozen hydrated fibrils exhibited one of two consistent helical crossovers. Fibrils with a 400Å helical crossover, which we termed the fast-twisting polymorph, accounted for only about 10% of fibrils present in the sample (Figure S3C). The remaining 90% of fibrils exhibited an 850Å helical crossover and were termed the slow-twisting polymorph (Figure S3D). The two polymorphs share very similar features apart from their different crossover distances, but given its abundance, the slow-twisting polymorph proved more amenable to high-resolution 2D classification and 3D structure determination. Subdivision of the fibril into two symmetry-related protofilaments could be inferred by the appearance of mirror symmetry in 2D class averages. Fibril images gave the impression of being mirrored across the fibril axis when it aligned parallel to the image plane. In this view, very near the fibril axis, the density of one protofilament appeared staggered relative to the other, resembling the teeth of a zipper (Figure S3E and S3F). These observations indicated that the two protofilaments might be related by a symmetry that approximated a 2_1_-screw axis. X-ray fibril diffraction from the LECT2 fibrils showed a strong 4.69Å reflection; corresponding to the true spacing between LECT2 molecules within a protofilament (Figure S2A and S2B). This spacing was confirmed by analysis of Fourier Transforms from high-resolution images of the fibrils (Figure S3B). The 2D classes were sufficient to generate a *de novo* 3D initial model in RELION and this map was used as an initial model for 3D classification (Figure S3G and S3H).

A three-dimensional reconstruction from the LECT2 fibril images displayed an ordered amyloid core with an estimated resolution of ~2.7Å, based on the 0.143 Fourier shell correlation (FSC) criterion (Figure S6B). The map was of sufficient quality to allow for building of an unambiguous atomic model. There were no breaks in connectivity of the tube of density attributed to each molecule and there was clear separation between layers of density corresponding to individual LECT2 molecules, whether stacked along the fibril axis or mating with the opposite protofilament (Figure 1A and 1B). Further, the resolution of the map allowed for the unambiguous assignment of side chains and peptide backbone oxygen atoms, allowing *de novo* assignment of the residues at its core (Figure 1C)^21,22^. The modelled portion of the amyloid core exhibited a clear pseudo-2_1_-screw symmetry with the repeating unit being a 21-residue sequence stretching from Methionine 55 to Isoleucine 75 (Figure 2A and 2B). To assure that the assignment of this sequence was correct, Rosetta was used to calculate configurations for all possible 21 residue segments of LECT2 constrained to match the observed density. This was done by threading each sequence onto the peptide backbone of the model to test which threading segment produces the most stable structure. This analysis confirmed that the M55-I75 sequence was the most energetically favorable sequence assignment for the geometry of the fibril core (Figure S7A).

**Figure 1:**
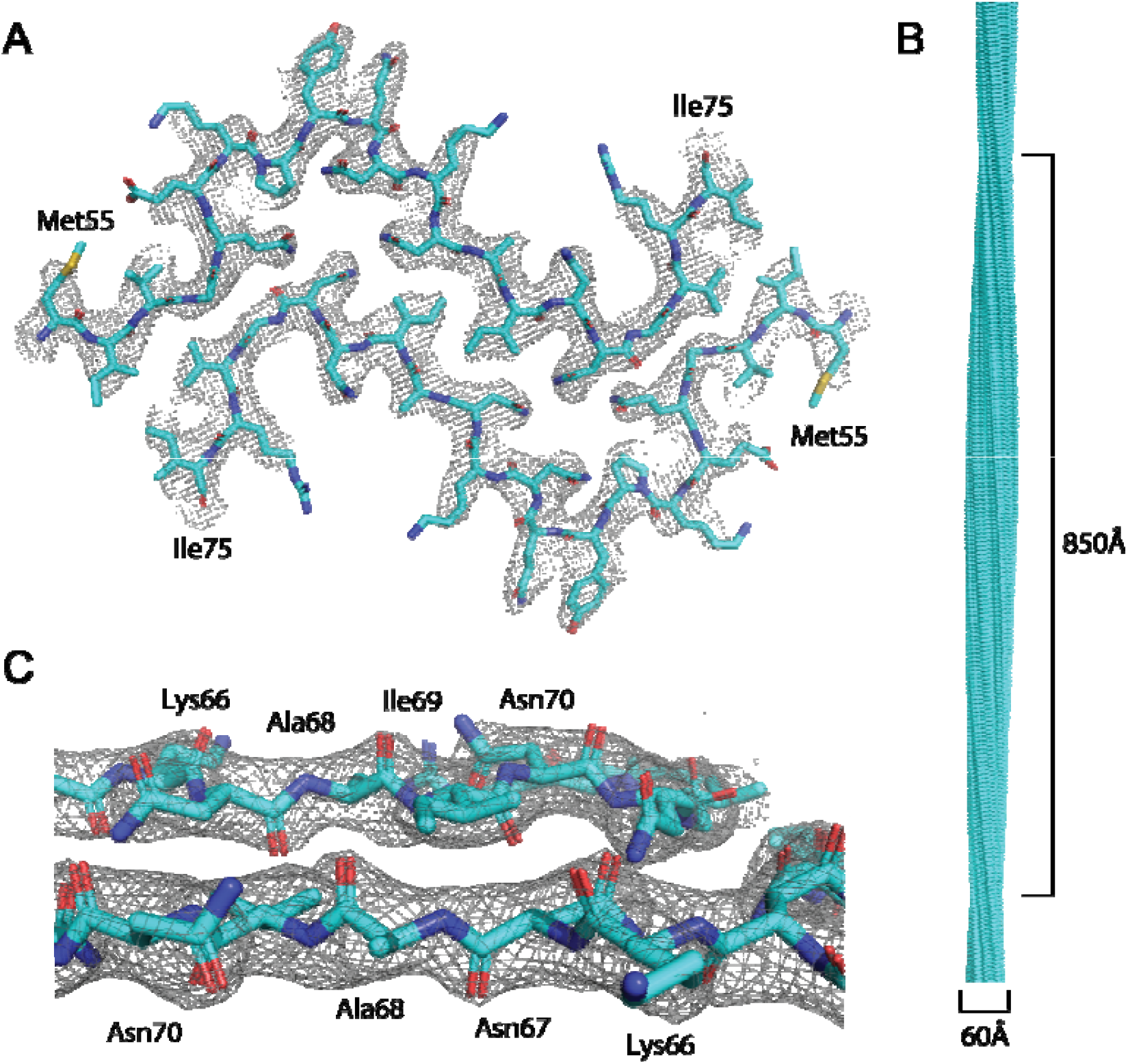
Views of the recombinant LECT2 fibril core revealed by cryo-EM helical reconstruction. A) Model fibril core structure of LECT2 (teal) modelled within the refined 2.7Å resolution cryo-EM map (grey mesh). B) View of the LECT2 fibril core with helical symmetry applied to show its full 850Å twist and 60Å width. C) Side view of the fibril model in the final cryo-EM map demonstrates its fit into density within the fibril layers.

**Figure 2:**
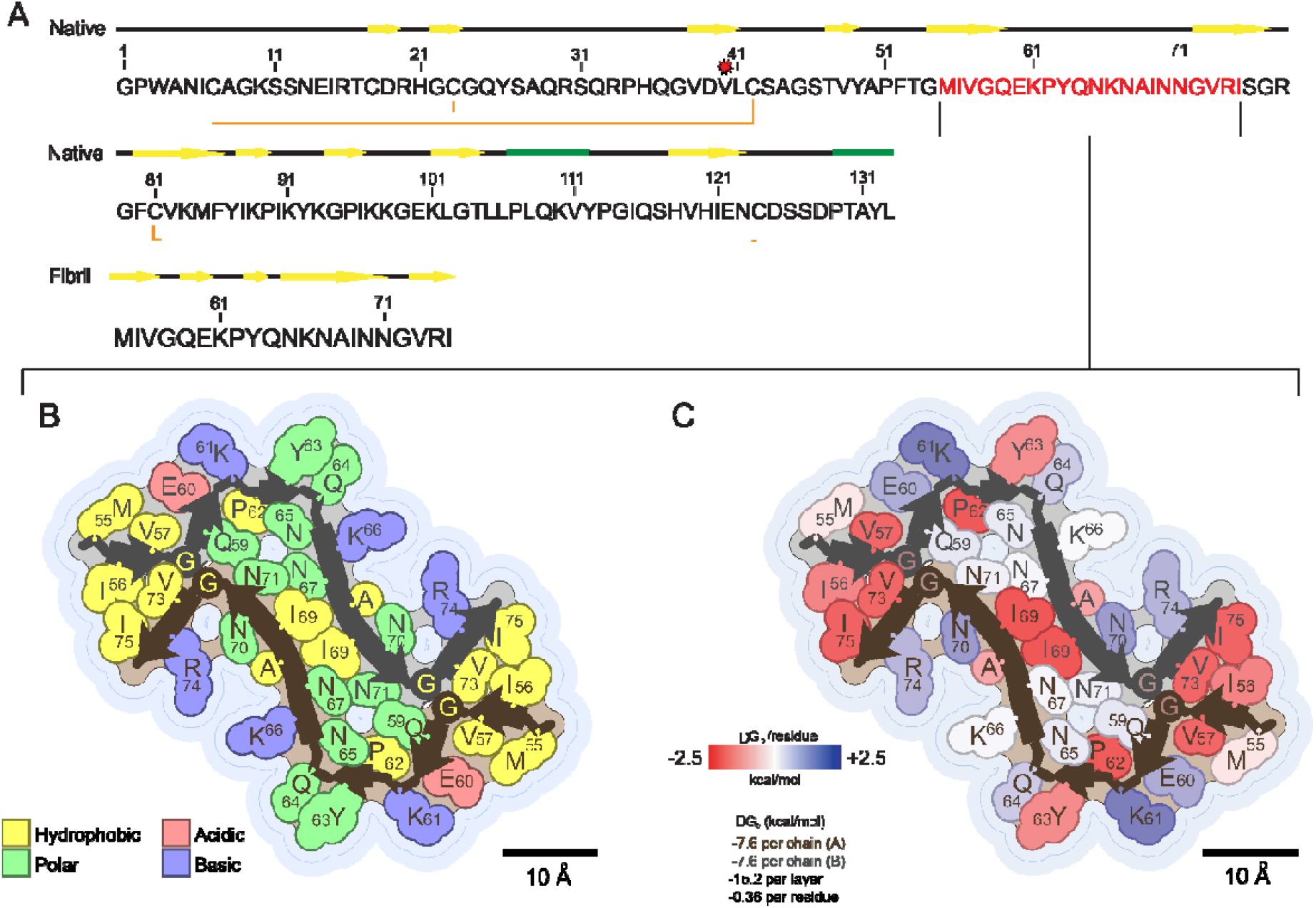
LECT2 fibril core sequence and structure overview. A) Sequence of LECT2 showing native secondary structure elements as yellow arrows for beta-strands and green rectangles for alpha helices with disulfides shown as orange brackets and the I40V polymorphism noted by a red star. The sequence of the fibril core is highlighted in red. B) Depiction of the fibril core showing the distribution of polar and nonpolar residues. C) Depiction of the fibril core showing stabilizing residues and the calculated solvation energy of the fibril core structure. Depictions of the fibril core were generated using the Amyloid Illustrator web service (https://srv.mbi.ucla.edu/AmyloidAtlas/Illustrator/).

### Stability-promoting molecular features in the LECT2 fibril core

At the core of the LECT2 fibril, a segment from each of two protofilaments mate in a largely dry interface. The two protofilaments are nearly 2-fold symmetric in the fibril core, allowing hydrophobic zippers to coalesce a network of uncharged polar residues stabilized by polar ladders. Each hydrophobic pocket within the fibril core contains tightly packed steric zipper structures composed of residues I56, V73, and I75; residue I69 anchors the very center of the fibril core near the helical axis (Figure 3A and 3B). A pair of glycines, G58 and G72, spaced only 4Å apart segregate the hydrophobic patches at the edges of the fibril core from the network of polar residues within it (Figure 3A). These polar residues include Q59, N65, N67, and N71, all of which are incorporated into stabilizing polar ladders (Figure 3C). Residues N67 and N71 form a polar clasp that holds together not only layers above and below but also stitches together the molecules from opposite protofilaments (Figure 3C). This polar clasp also seals off the central pair of I69 residues and pins together the two largest beta-strands in the fibril structure. That combination of features results in an overall energetically favorable structure with a calculated standard free energy of solvation^2,23^ of ΔG_o_ of -15.2 kcal/mol per layer and -0.36 kcal/mol per residue (Figure 2C). The solvation energy and small size of the LECT2 amyloid core are similar to that of the human RIPK1-RIPK3 hetero-amyloid, which is hypothesized to form stable aggregates *in vivo* and signal TNF-induced necroptosis in cells^24^. These metrics could be further influenced by other stabilizing interactions, such as disulfide bridges or metal coordination sites, that in the present structure are absent from the fibril core, but exist within the fibril fuzzy coat and remain unresolved.

**Figure 3:**
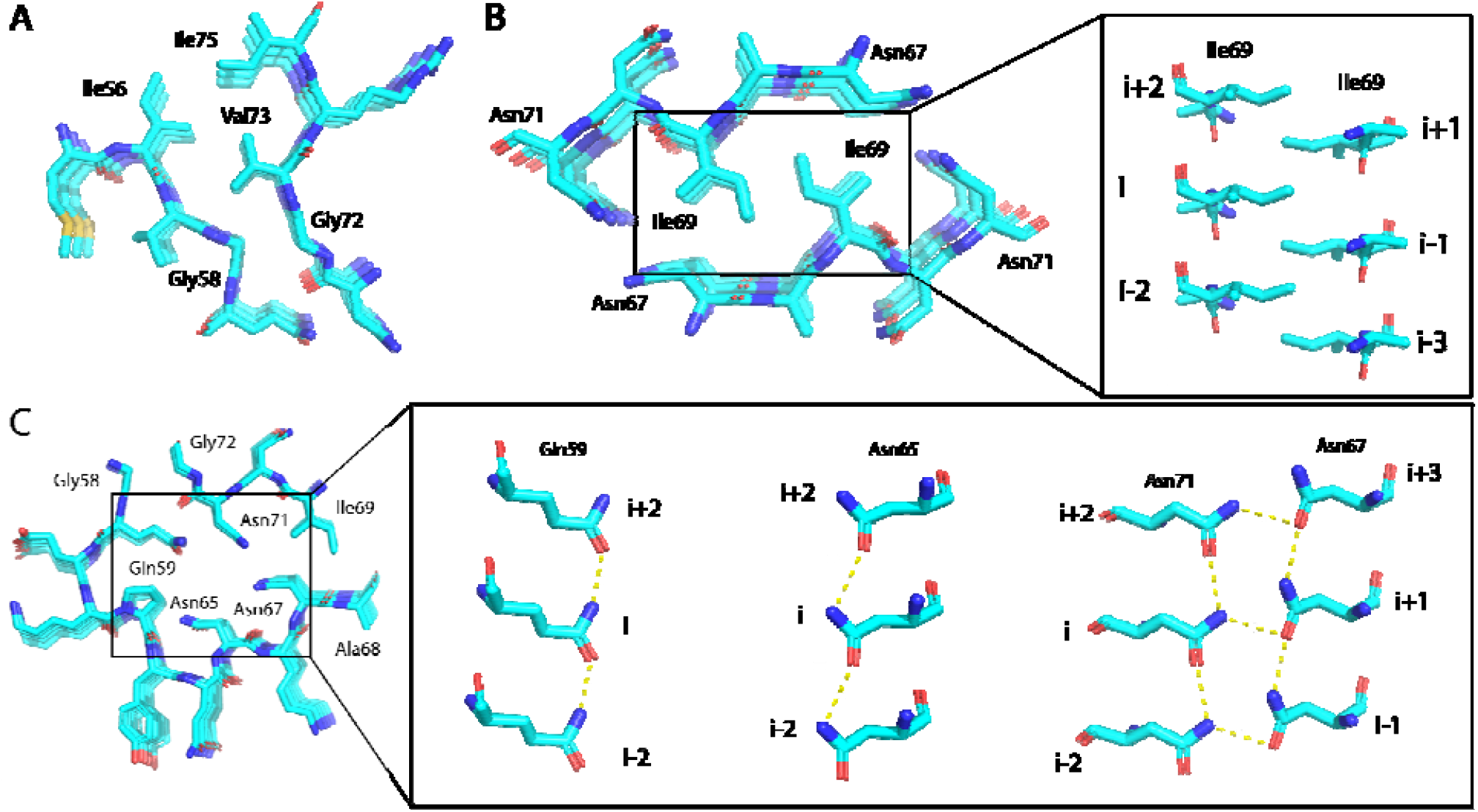
Stabilizing contacts at the core of the LECT2 fibril. A) The outer steric zipper interface formed by residues Gly58, Gly72, Val73, Ile56, and Ile75. B) The central steric zipper interface formed by residue Ile69 is flanked by a paired polar ladder. Inset shows the pseudo-2_1_ stacking of fibril layers. C) Hydrophilic pocket within the LECT2 fibril core contains polar ladders formed by uncharged polar residues Gln59, Asn65, and Asn67 with Asn71, spaced 4.69Å apart along the fibril axis (inset).

## Discussion

In search of amyloid folds adopted by the LECT2 protein, we find that recombinant LECT2 polypeptides encoding the I40V sequence polymorphism readily form amyloid fibrils that display a canonical beta-sheet rich architecture with stacked layers of the LECT2 protein separated by a rise of 4.69 Å per layer and obeying a regular, left-handed, helical twist. The amyloid core resolved within these fibrils is composed of two identical protofilaments that each contain 21 of the 133 residues in full-length LECT2: from Methionine 55 to Isoleucine 75. Interestingly, this sequence has not been predicted by previous work to form amyloid fibrils. However, other studies identified the regions immediately preceding and following it as possible amyloidogenic segments^20^, and the newly identified core segment is predicted to form steric zipper structures by the amyloid-prediction algorithm in ZipperDB^25^(Figure S1A).

While a significant fraction of the LECT2 sequence is predicted to be amyloidogenic, much of the protein’s globular fold is stabilized by three disulfide bonds, two in the N-terminal region and one in C-terminal region (Figure 2A). These bonds would likely prevent many of the predicted amyloidogenic regions from assembling into fibrils. However, the core segment identified here is not involved in any of the native disulfide bonds and represents a portion of the central beta-barrel motif of LECT2^17^. Specifically, the LECT2 fibril core encompasses most of beta-strands 3 and 4 and all of loop 2 in the LECT2 globular fold (Figure S8A). That loop is solvent-facing and sits adjacent to the Zn(II) binding pocket^17^. This same region of the protein has previously been hypothesized to be susceptible to reductions in pH that could facilitate the loss of the bound zinc coordinated to the first and second beta-strands and may destabilize the beta-barrel structure, leading to the exposure of amyloidogenic segments^18^.

A comparison of native and fibrillar LECT2 structures reveals that the beta-strand conformation of N- and C-terminal segments of the fibrillar fold are retained from beta-strands 3 and 4 of the globular fold. However, the central segment of the fibril core represents a large ordering of secondary structure, namely rearrangement of loop 2 into 3 beta-strand segments, the largest of which stretches from N65 to N71 (Figure 2B). The outer surface of the LECT2 fibril is decorated with larger amino acids including basic residues K61, K66, and R74 as well as one acidic residue, E60, and a pi-stacked Y63 (Figure 2B). These residues are also in solvent-exposed positions in the globular fold of LECT2. Residues I56, V73, and I75, which form the outer steric zipper segments of the fibril, are all buried within the central beta-barrel of globular LECT2, and I69, the central hydrophobic residue, is natively buried as part of loop 2 abutted to alpha-helix 1 (Figure S8B). In contrast, the polar-ladder-forming residues Q59, N65, N67, and N71 undergo a large conformational and environmental change from the globular fold. All four of these residues are solvent-facing in loop 2 of the globular structure of LECT2, and all are packed into a relatively dry interface in the fibril structure (Figure S8B). Polar ladder conformations like those revealed in this structure have long been hypothesized to stabilize amyloid assemblies^26,27^ and were also favored during energy minimization of the LECT2 fibril fold in Rosetta (Figure S7B). The structural transition to a polar ladder within a dry environment confers increased stability by joining chains along the fibril axis while fully satisfying hydrogen bonds within the fibril core. This stabilizing effect is further amplified when the two polar ladders in separate protofilaments interdigitate into a zipper conformation, as is observed between residues N67 and N71 in the LECT2 fibril.

It remains to be seen whether recombinant LECT2 amyloid fibrils exactly match those present in the kidneys of ALECT2 patients; however, structures of the former and the segments at their core provide a first glimpse at stabilizing interactions within LECT2 fibrils. Further, the exact role of the I40V polymorphism in amyloid aggregation remains unclear as the residue was not resolved as part of the LECT2 amyloid core. The segment at the core of this recombinant LECT2 fibril is derived from part of the central beta-barrel structure of globular LECT2, supporting the theory that loss of zinc may destabilize this portion of the protein, exposing the amyloidogenic segments resolved here and allowing for aggregation. This information opens the possibility for rational design of aggregation inhibitors which could either stabilize this region of globular LECT2 or inhibit the extension of fibrils in the amyloid form of LECT2 by targeting residues at its core.

## Methods

### Protein Expression and Purification

Full-length, mature human sequence LECT2 protein encoding the I40V sequence polymorphism and an N-terminal 6-His tag was transformed into and expressed in BL21-gold E. coli cells. Two 1L cultures grown at 37°C to an O.D. of 0.75 were induced with 1mM IPTG and allowed to shake overnight at 23°C. Cells were then collected by centrifugation at 10,000xg for 10 minutes, resuspended in 1X PBS with 6M guanidine, and lysed on an Avestin Emulsiflex C3 (ATA Scientific). Insoluble cell material was removed by centrifugation at 9000 rpm for 50 minutes and the supernatant was collected, filtered using a 0.8µm size cutoff, and run over a His-trap column and washed with 3 column volumes of buffer. Protein was eluted with an elution buffer of 6M guanidine, 1X PBS, and 500 mM imidazole. Eluate fractions were evaluated for protein purity by gel electrophoresis then pooled and concentrated to 40 mg/ml (Figure S1B). Concentrated stocks were flash frozen for storage at -80°C in 1X PBS with 6M guanidine and 20mM TCEP.

### Thioflavin T Fluorescence

To generate a Thioflavin T (ThT) fibril growth kinetics curve, LECT2 protein was prepared to 2mg/ml in fibrillization buffer as described above. Equimolar amounts of Thioflavin T, also dissolved in fibrillization buffer, was added to this solution. This preparation along with a control of only fibrillization buffer and ThT were set up in triplicates of 100 µL in a 96 well plate. The samples were then subjected to continuous shaking on a Varioskan LUX (ThermoFisher Scientific) as the ThT fluorescence was measured every 15 minutes for 48 hours using an excitation wavelength of 445 nm and measuring emission at 482 nm.

### Cryo-EM Sample Preparation and Data Collection

To grow fibrils for cryo-EM analysis, stock LECT2 protein was diluted to 2 mg/ml (120 µM) into a fibrillization buffer containing 50 mM MOPS 6.5, 30 mM TCEP, 10 mM EDTA, and 150 mM NaCl. A volume of 100 µL of protein solution per well was then agitated in a 96 well plate on an acoustic shaker running at 60 Hz (2400 rpm) at 37°C for 48 hours. The solution was collected and spun down at 10,000xg for 5 minutes to remove large aggregates not amenable to single particle analysis. The solution was then diluted 6-fold in buffer and glycerol was added to the remaining supernatant to a concentration of 0.25%. A volume of 1.5 µL of this solution was added to each side of a gold Quantifoil R 1.2/1.3 cryo-EM grid (Ted Pella Inc.) within a Vitrobot System (ThermoFisher Scientific) with humidity set to 100% and a temperature of 4°C. The grid was then blotted for 1.5 seconds with the blot force set to -1, plunge frozen in liquid ethane, and stored for data collection in liquid nitrogen. High-resolution cryo-EM data was collected at the Stanford-SLAC Cryo-EM Center (S^2^C^2^) over the course of 48 hours using EPU for automated data collection. A total of 13830 movies were obtained taking three shots per hole on the grid. Movies were collected on a Titan Krios G3i microscope (ThermoFisher Scientific) equipped with a BioQuantum K3 camera (Gatan, Inc.) using a pixel size of 0.79□Å/pixel, a total dose of 52□e−/Å^2^ over 40 frames, and a defocus range between −0.8 and −1.8□μm.

### Cryo-EM Data Processing

Collected images were input into MotionCor2 for drift correction and CTFFIND4 was used to calculate Contrast Transfer Functions (CTF). Fibrils were automatically picked using filament mode in crYOLO^28,29^ after being trained on a manually selected pool of fibril segments picked from 100 movies. This yielded 1,622,942 segments using an overlap of 90% between neighboring segments and a box size of 384 pixels. The segments were then transferred to RELION 3.1^30–32^ for all subsequence 2D and 3D classification. The segments were split into six equally sized groups and subjected to iterative rounds of reference-free two-dimensional (2D) classification using T□=□8 and K□=□100 to remove poorly aligned classes. After narrowing each split of segments, they were combined again (TL=L8 and KL=L100) resulting in 308,578 segments contributing to well-defined 2D classes to be used in 3D classification. These classes were used to generate a *de novo* 3D initial model which was used as a reference for further 3D classification. Initial classification (K = 4, T = 4) did not impose pseudo-2_1_-screw symmetry and used an initial helical twist of -1.06° and a helical rise of 4.8 Å. The best class from this classification was selected out and further subclassified by removing any segments with a CTF estimate poorer than 4 Å resolution. The resulting 51,543 segments were used for further 3D classification (K = 3, T = 4) allowing local optimization of helical twist and rise. The best resulting class showed clear strand separation along the fibril axis and the pseudo-2_1_-screw symmetry became apparent without it having been imposed. The model and data from these 24,770 segments were used for high-resolution gold-standard 3D auto-refinement with the pseudo-2_1_-screw symmetry now enforced. The resulting model underwent iterative Bayesian polishing^32^ and CTF refinement^33^ to further improve the map (Figure S6A). A final refined helical twist□of□179.49° was derived from the auto-refine map and a helical rise of 2.345 Å was imposed on the post-processed map based on X-ray fibril diffraction data showing 4.69 Å strand separation in the fibril samples. The final resolution was ultimately calculated to be 2.715 Å from gold-standard Fourier shell correlations at 0.143 between two independently refined half-maps (Figure S6B).

### Model Building

A poly-alanine model was initially built into the post-processed map in Coot^34^. Side chains were modified manually to test the fit of different registrations of the LECT2 sequence in the density. A close approach between tubes of density could be interpreted only as the G58-G72 interaction between symmetric strands, as the closeness of the approach would sterically exclude all other side chains. This feature helped to identify the likely segment of LECT2 and the entire 21 residue chain from Met55-Ile75 was built out from there. Final atomic refinements and statistical calculations for Supplementary Table 1 were performed in Phenix^35^. While the map seemed to agree clearly with this assignment, we decided to confirm its validity by testing the fit and favorability of all possible 21-residue segments from the LECT2 sequence within the map using Rosetta. Threading was performed with a custom python script utilizing the PyRosetta software package^36^. In a sliding window approach, each sequence window was threaded onto a poly-alanine backbone, placed in the density and Fast Relaxed in PyRosetta. The pipeline used the REF2015 score function^37^ with an increased electrostatics weight (fa_elec = 1.5) in combination with a density score term (elec_dens_fast = 25). Each sequence was energy minimized in triplicate and the resulting scores were averaged. Symmetry was applied to pose objects to increase speed of computation. The Fast Relaxed structure was then optimized and refined in Coot and Phenix^35^.

### X-ray Fibril Diffraction

LECT2 amyloid fibrils were grown as described above for cryo-EM sample preparation. A 3 uL droplet of the mature fibril solution was pipetted between two glass rods held about a millimeter apart and allowed to evaporate. This process was repeated multiple times until a visible fibril bundle was observed bridging the glass rods. This bundle was place on an in-house Rigaku FRE+ rotating anode X-ray beamline and exposed for 5 minutes onto a Rigaku HTC detector with varimax confocal optics. Proteinase K and ice ring diffraction patterns were used to calibrate the detector distance and accurately measure the 4.69 Å reflection observed from the fibril bundle.

### Fibril Stability Assays

LECT2 amyloid fibrils were grown as described above for cryo-EM sample preparation and then pelleted at 14,300xg for 10 minutes. The fibril pellets were then resuspending to a protein concentration of 40 µM in water, fibrillization buffer, 3M urea with fibrillization buffer, and fibrillization buffer with 1:1000 Proteinase K (Sigma-Aldrich) to a final concentration of 40 nM. These solutions were immediately pipetted in triplicate into a 96 well tray alongside buffer control wells for each and placed on a NEPHELOstar Plus plate reader (BMG Labtech). The tray was stirred every ten minutes to homogenize the mixture and light scattering of particles in solution was measured immediately after each agitation. For analysis of protease resistant LECT2 fragments, fibrils at 120 μM were incubated with 10 μM Proteinase K for various timepoints and then evaluated by SDS-PAGE (Figure S9A). The protein gel band from the one-hour timepoint was extracted, digested with Trypsin, and analyzed by gel liquid chromatography tandem mass spectrometry.

### In-gel digestion and peptide mass fingerprinting of LECT2 using GeLC-MS/MS

Gel Liquid Chromatography tandem mass spectrometry spectra collected on Proteinase K digested LECT2 fibrils were acquired on a ThermoFisher Q-Exactive Plus (UCLA Molecular Instrumentation Center, Los Angeles, CA, USA). LECT2 fibrils at 120 μM were incubated with 10 μM Proteinase K for one hour. These samples were removed at various timepoints, boiled for 10Lminutes at 98□°C to halt digestion, and loaded on a 4–12% Bis-Tris SDS-PAGE gel. The gel band from the one hour timepoint was excised and digested with 200 ng trypsin at 37°C overnight. The digested products were then extracted from the gel bands in 50% acetonitrile/49.9% H2O/ 0.1% trifluoroacetic acid (TFA) followed by desalting with C18 StageTips. Extracted peptides were then injected on an EASY-Spray HPLC column (25□cm□×□75□µm ID, PepMap RSLC C18, 2□µm, ThermoScientific) and tandem mass spectra were acquired with a quadrupole orbitrap mass spectrometer (Q-Exactive Plus Thermo Fisher Scientific) interfaced to a nanoelectrospray ionization source. The raw MS/MS data were converted into MGF format by Thermo Proteome Discoverer (VER. 1.4, Thermo Scientific and analyzed by a MASCOT sequence database search.

## Acknowledgements

This work is supported by DOE Grant DE-FC02-02ER63421, NSF Grant DMR-1548924, the NIH-NIGMS Grant R35 GM128867. L.S.R. is supported by the USPHS National Research Service Award 5T32GM008496. J.A.R. is also supported as a Sloan Fellow, a Pew Scholar, and a Packard Fellow. Part of this work was performed at the Stanford-SLAC Cryo-EM Center (S2C2). The content is solely the responsibility of the authors and does not necessarily represent the official views of the National Institutes of Health. We thank the following S2C2 personnel for their invaluable support and assistance: Dr. Patrick Mitchell, Dr. Yee-Ting Lee, and Professor Wah Chiu who are supported by the National Institutes of Health Common Fund Transformative High-Resolution Cryo-Electron Microscopy program (U24 GM129541). In addition, we thank the staff of the UCLA Molecular Instrumentation Center (MIC), supported by the UCLA Division of Physical Sciences.

## Data Availability

The reconstructed cryo-EM map was deposited in the Electron Microscopy Data Bank with the accession code EMD-29682, while coordinates for the refined atomic model were deposited in the Protein Data Bank under the accession code 8G2V.

## Supplementary Information

**Supplementary Table 1:** Cryo-EM data collection and structure information.

| **Data Collection** |  |
| --- | --- |
| Magnification | 110,000x |
| Defocus Range (um) | -0.8 to -1.8 |
| Voltage (kV) | 300 |
| Microscope | Titan Krios G3i |
| Camera | BioQuantum K3 camera |
| Frame exposure time (s) | 2.5 |
| No. of movie frames | 40 |
| Total electron dose (e<sup>-</sup>/ Å<sup>2</sup>) | 52 |
| Pixel Size (Å) | 0.799 |
| **Reconstruction** |  |
| Box size (pixel) | 384 |
| Inter-box distance (Å) | 30 |
| No. of segments extracted | 1,622,942 |
| No. of segments after Class2D | 308,578 |
| No. of segments after Class3D | 24,770 |
| Map Resolution (Å) | 2.715 |
| FSC threshold | 0.143 |
| Map sharpening B-factor (Å<sup>2</sup>) | -69.7 |
| Helical rise (Å) | 2.345 |
| Helical twist (°) | 179.49 |
| Symmetry imposed | C1 |
| **Atomic Model** |  |
| Initial model used | *De novo* |
| No. of unique non-hydrogen atoms | 166 |
| R.m.s.d. bonds (Å) | 0.006 |
| R.m.s.d. angles (°) | 1.055 |
| Molprobity score | 1.88 |
| Favored rotamers (%) | 94.74 |
| Ramachandran favoured (%) | 5.26 |
| Ramachandran outliers (%) | 0 |
| CB deviations > 0.25 Å (%) | 0 |
| Bad bonds (%) | 0 |
| Bad angles (%) | 0 |

**Supplementary Figure 1:**
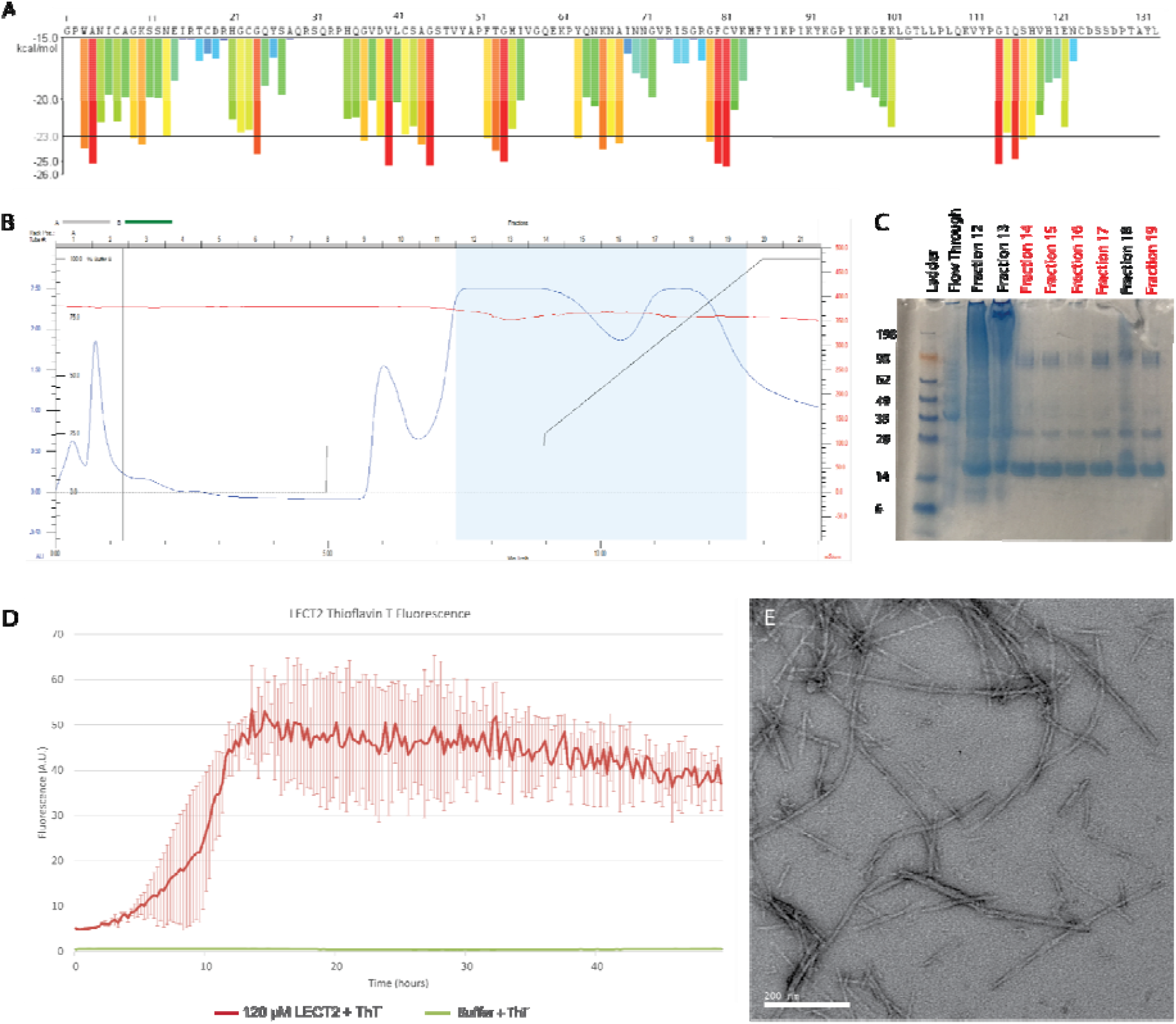
Production of recombinant LECT2 amyloid fibrils. A) Predicted amyloid forming six-residue segments (kcal/mol < -23.0) are shown for the LECT2 sequence, calculated using ZipperDB. B) A His-tagged, full-length LECT2 polypeptide was expressed and purified using a His-Trap column under the illustrated gradient. C) Purity of the fractions was evaluated by gel electrophoresis. The purest fractions: 14-17, 19 (red) were pooled and concentrated. D) The recombinant protein was used to form fibrils and Thioflavin T fluorescence was monitored over 48 hours of growth. E) The resulting fibrils from Thioflavin T analysis were imaged by negative stain TEM.

**Supplementary Figure 2:**
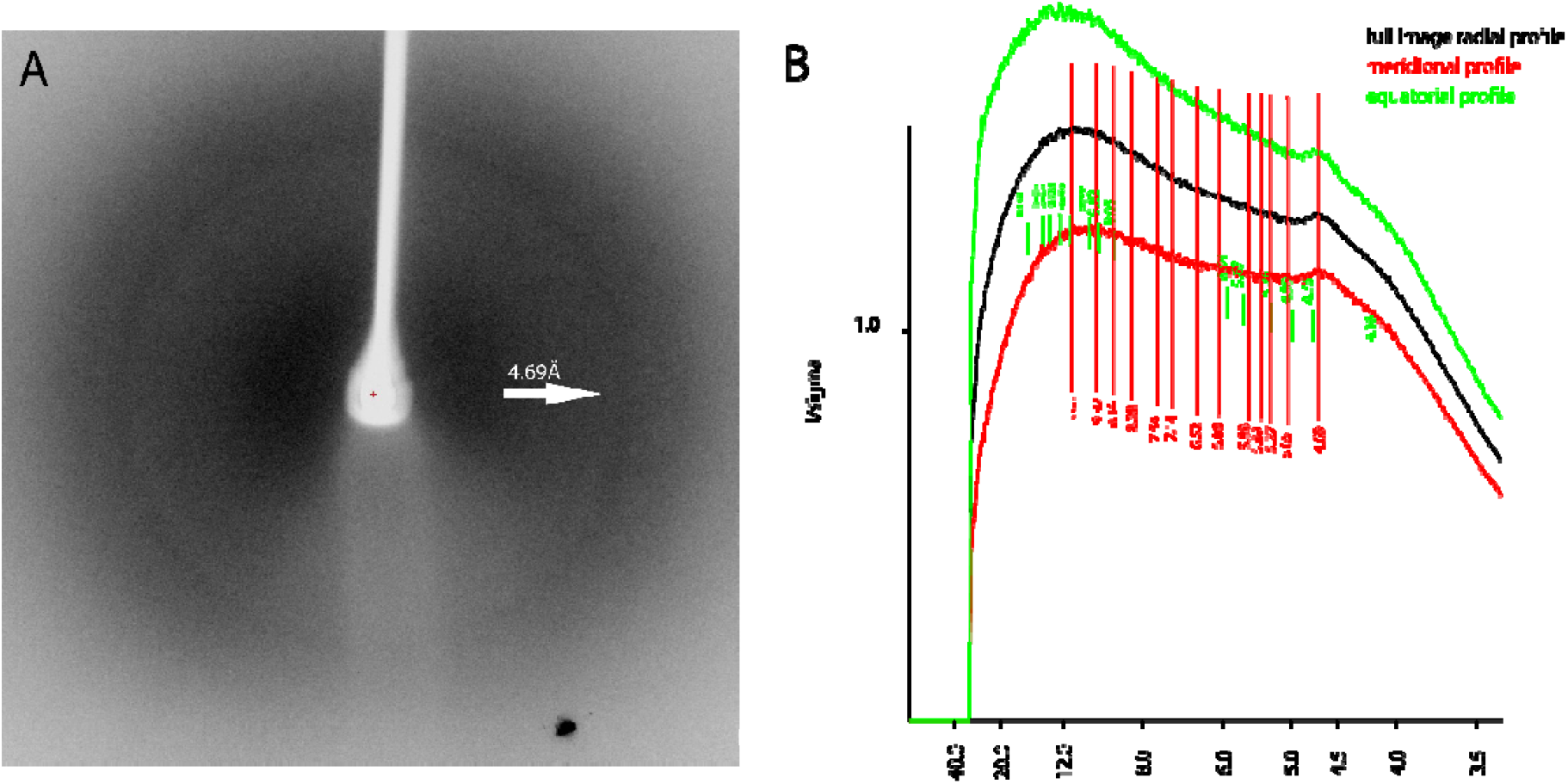
X-ray Diffraction from LECT2 fibrils A) X-ray diffraction exposure of bundles recombinant LECT2 fibrils. B) Radial profile generated from the diffraction image indicates a strong diffraction ring at 4.69Å.

**Supplementary Figure 3:**
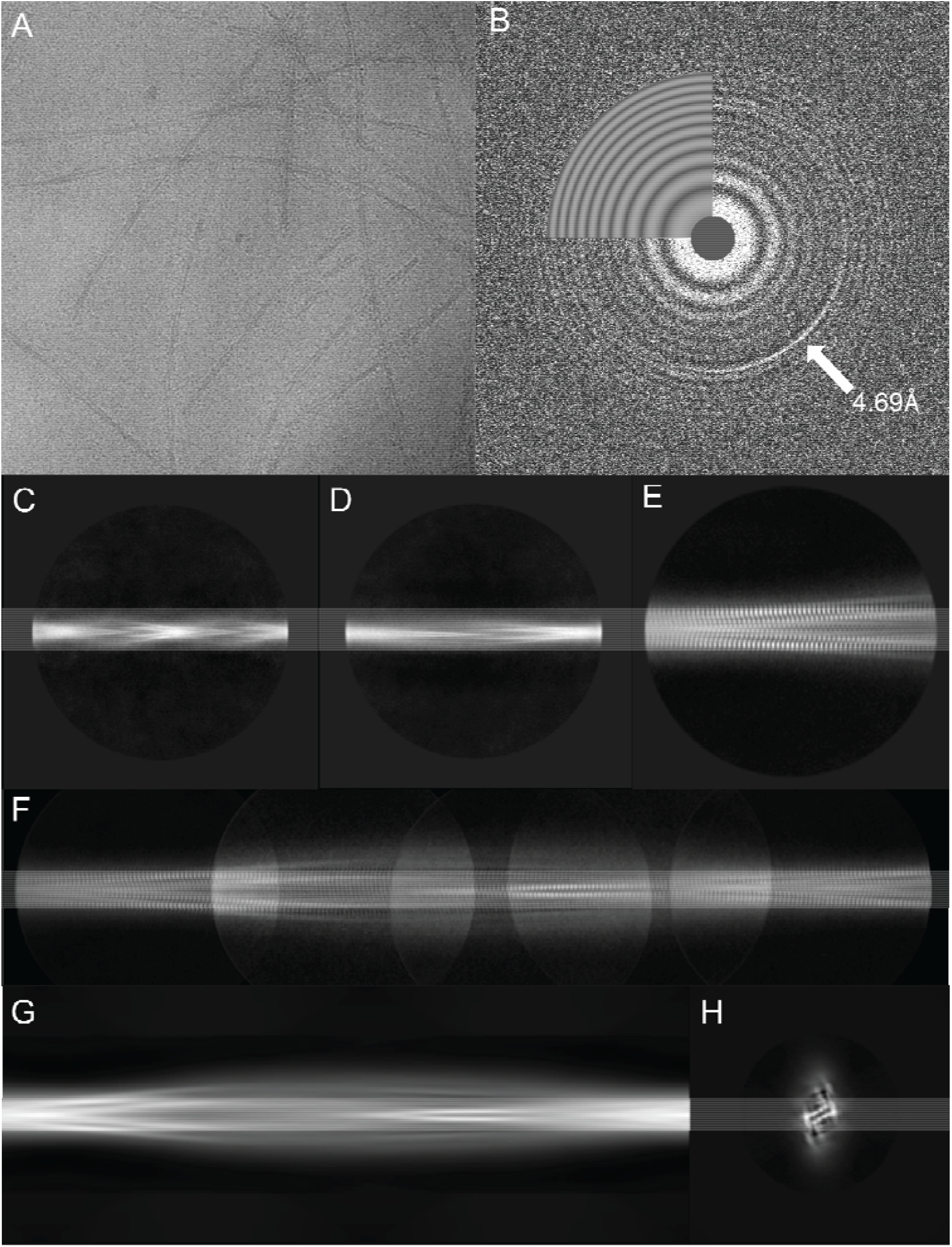
Vitrified LECT2 fibrils, CTF analysis, and 2D classification. A) Example image of the vitrified fibrils used for cryo-EM data collection. B) Example CTF image showing the signature fibril diffraction profile at 4.69Å. C) Representative 2D class of the fast-twisting fibril species from a 1024 pixel box. D) Representative 2D class of the slow-twisting fibril species from a 1024 pixel box. E) A slow-twisting fibril image from 2D class after classification using a 384 pixel box in Relion. F) Composite image of 384 pixel box size 2D classes from the slow-twisting fibril polymorph stitched together to show its full twist. G) *De novo* 3D model generated from 2D classes in Relion viewed from the side and H) as a cross section.

**Supplementary Figure 4:**
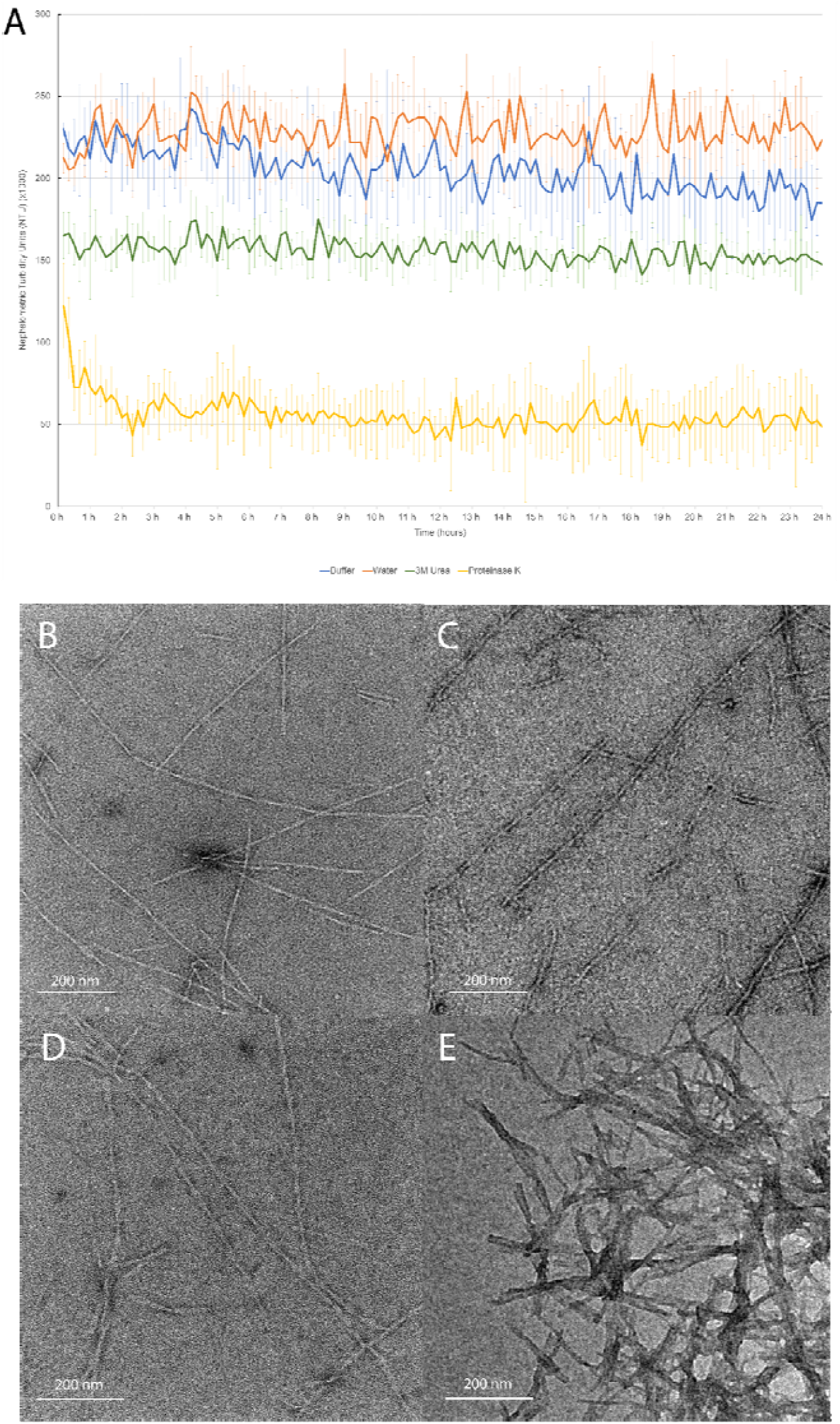
LECT2 fibril stability. A) Nephelometer turbidity readings for the incubated fibril samples over 24 hours. B-E) Negative stain TEM images of the fibrils remaining after 24 hours incubation in fibrillization buffer (B), water (C), 3M urea (D), and 40nM Proteinase K (E).

**Supplementary Figure 5:**
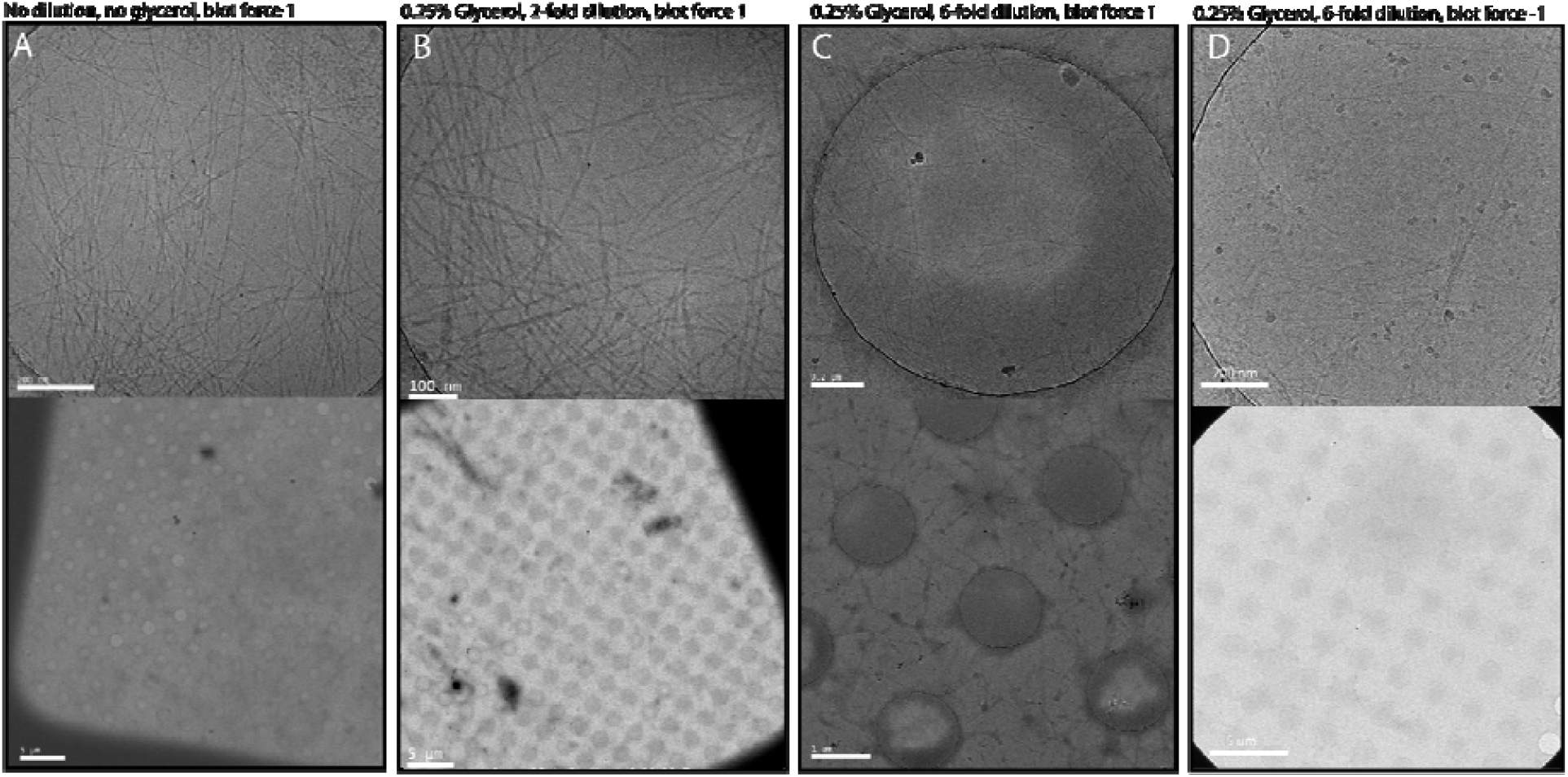
Cryo-EM grid optimization. Recombinant fibrils were plunge frozen in buffer at different dilutions and Vitrobot blot settings for optimal particle dispersion, representative micrographs are shown. A) No dilution results in clumping and overcrowding. B) Addition of 0.25% glycerol improves ice distribution, but 2-fold dilution does not reduce crowding. C) 6-fold dilution improves fibril distribution in grid holes. D) Reducing the blot force improves ice distribution and maintains fibril distribution.

**Supplementary Figure 6:**
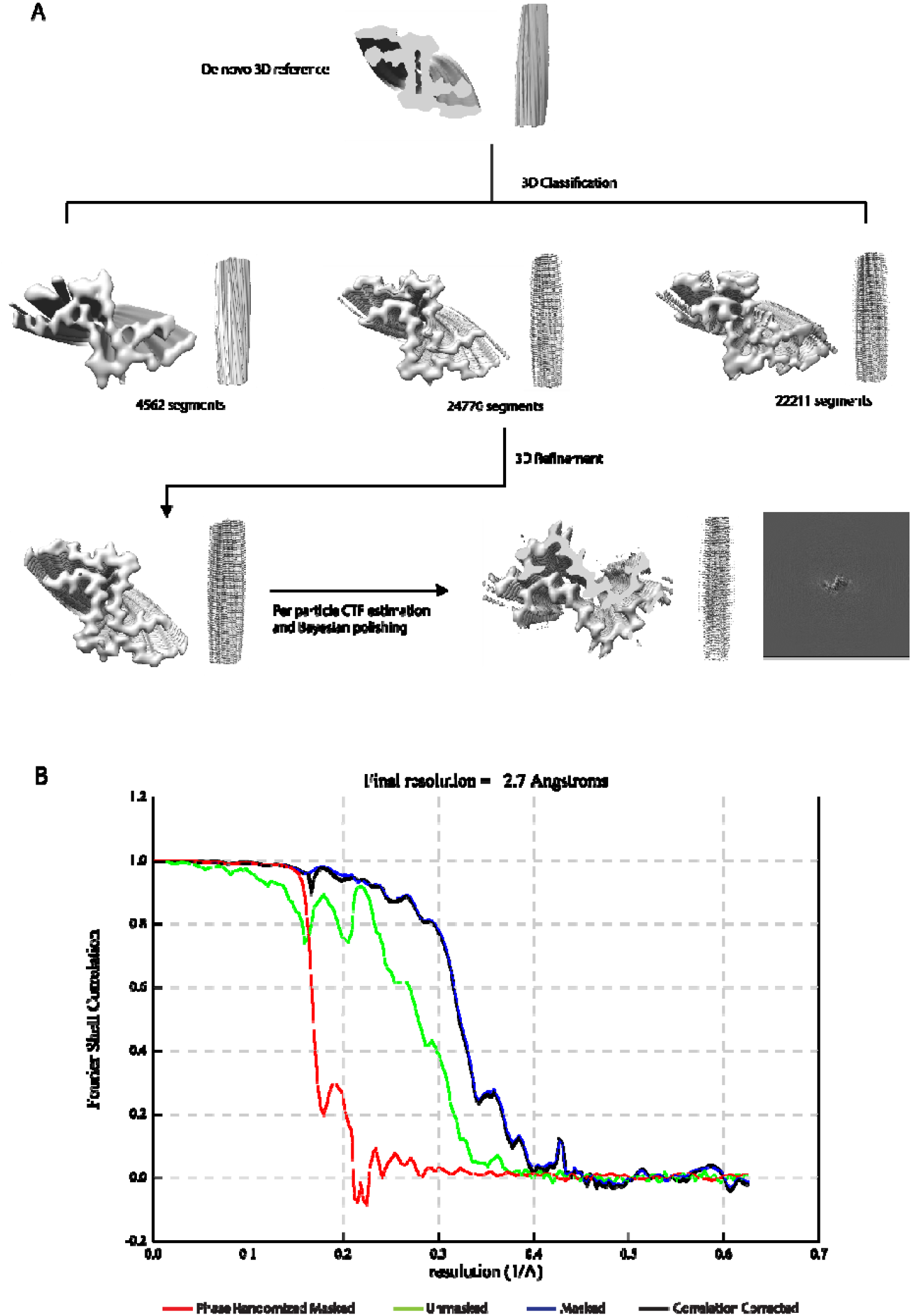
Cryo-EM data processing and model comparisons. A) Overview schematic representation of helical reconstruction of fibril species performed in RELION showing maps of fibril sides and cross sections. B) FSC curves between independently refined half-maps which were phase randomized (red), unmasked and corrected (green), masked (navy blue), or masked and corrected (black). Final resolution of 2.7Å was calculated using a cutoff of FSC=0.143.

**Supplementary Figure 7:**
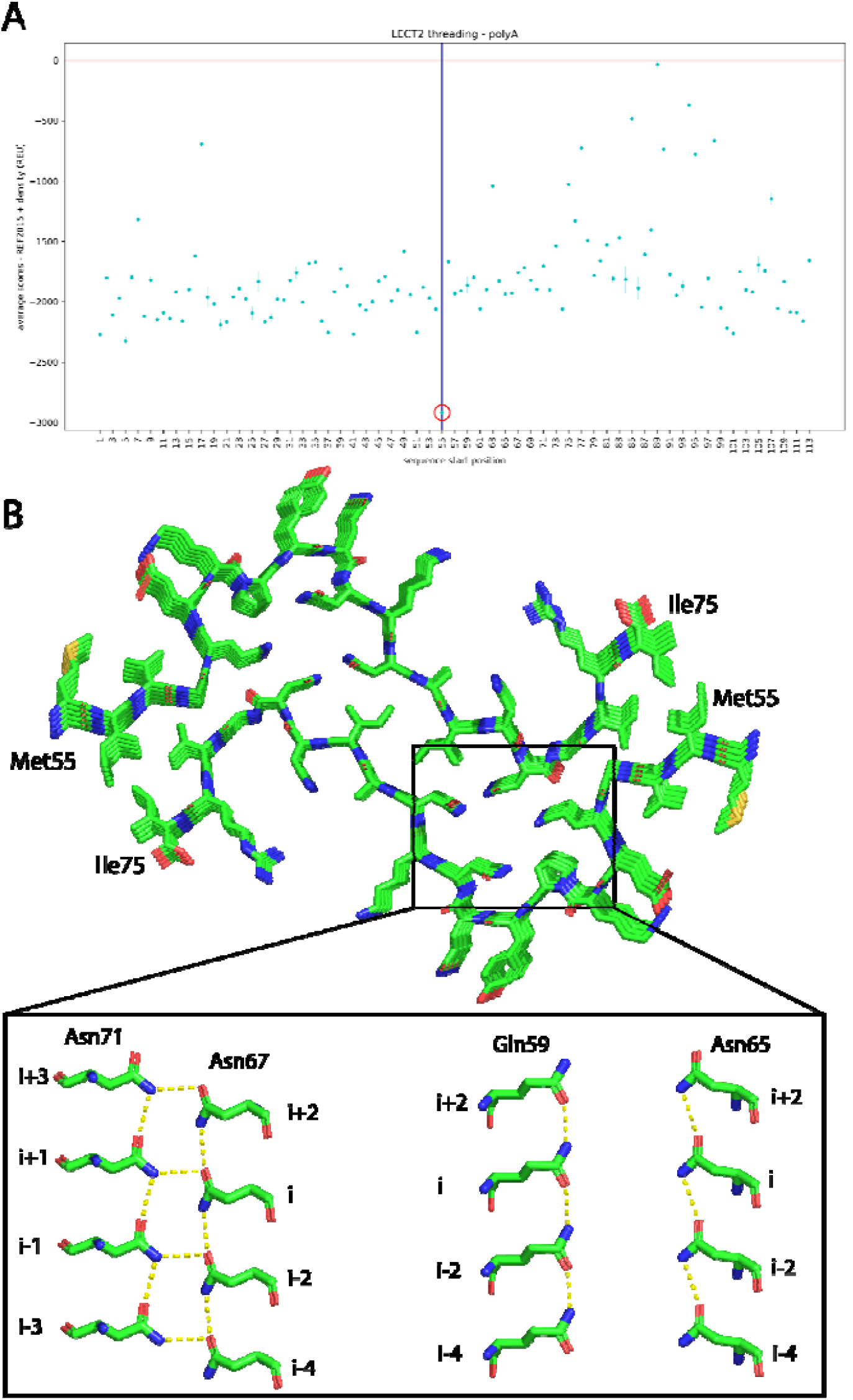
Identification of sequence matching LECT2 fibril structure by threading and energy minimization in Rosetta. A) Threading analysis indicates that the sequence beginning at Met55 is the most energetically favorable among possible assignments for the cryo-EM map (red circle). B) Energy-minimized, fast relaxed model from Rosetta predicts overall fibril core morphology as well as the stabilizing polar ladders in the hydrophilic pockets (inset).

**Supplementary Figure 8:**
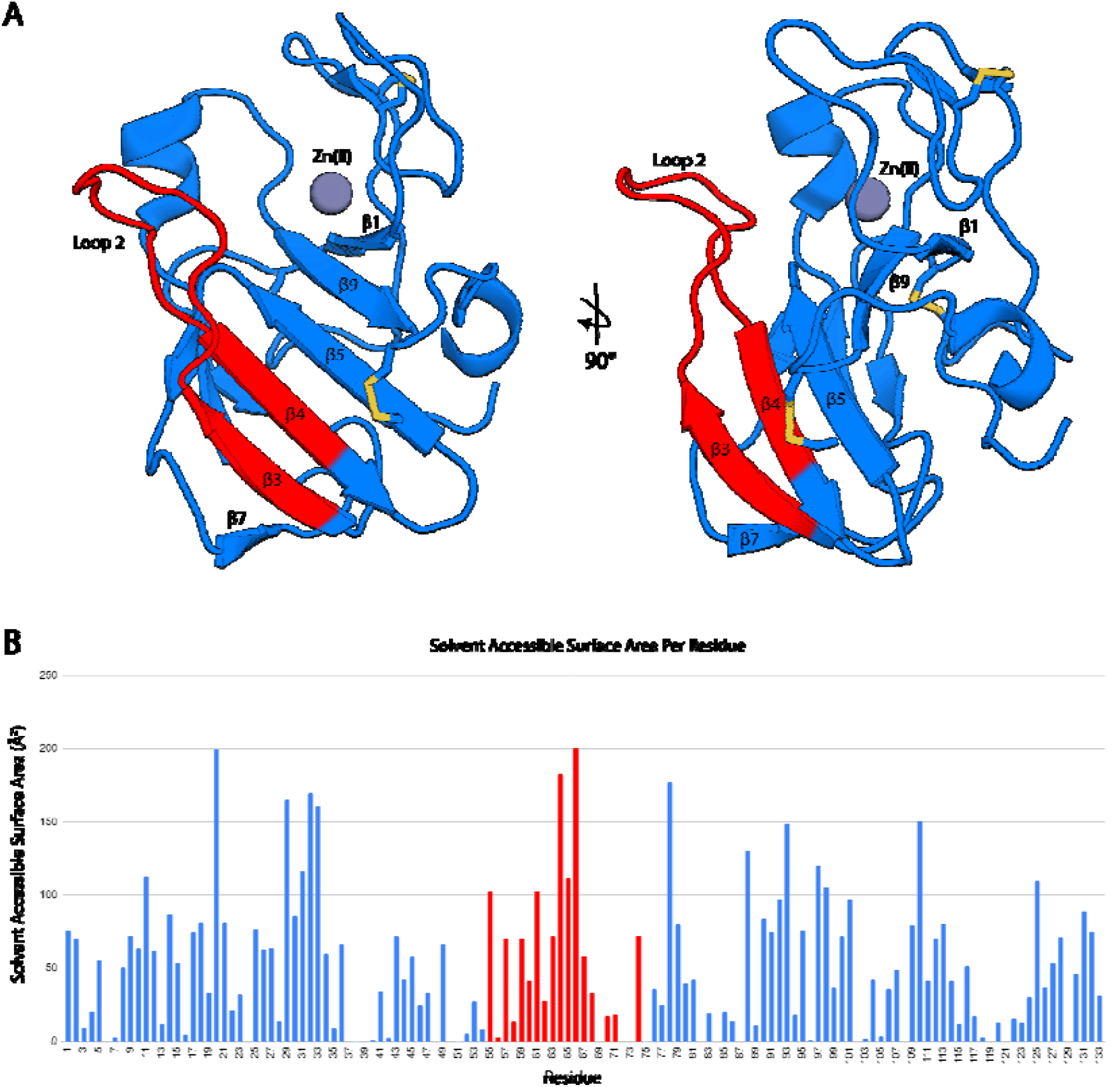
LECT2 globular fold and solvent accessibility of residues 55-75. A) Views of LECT2 in its globular form (PDB:5B0H)^17^. Residues 55-75 (colored in red) are identified as forming the amyloid core of LECT2 in its fibril form. B) Solvent accessible surface area per residue of LECT2 in its globular fold calculated using the Ochanomizu University program (ver. 1.2).

**Supplementary Figure 9:**
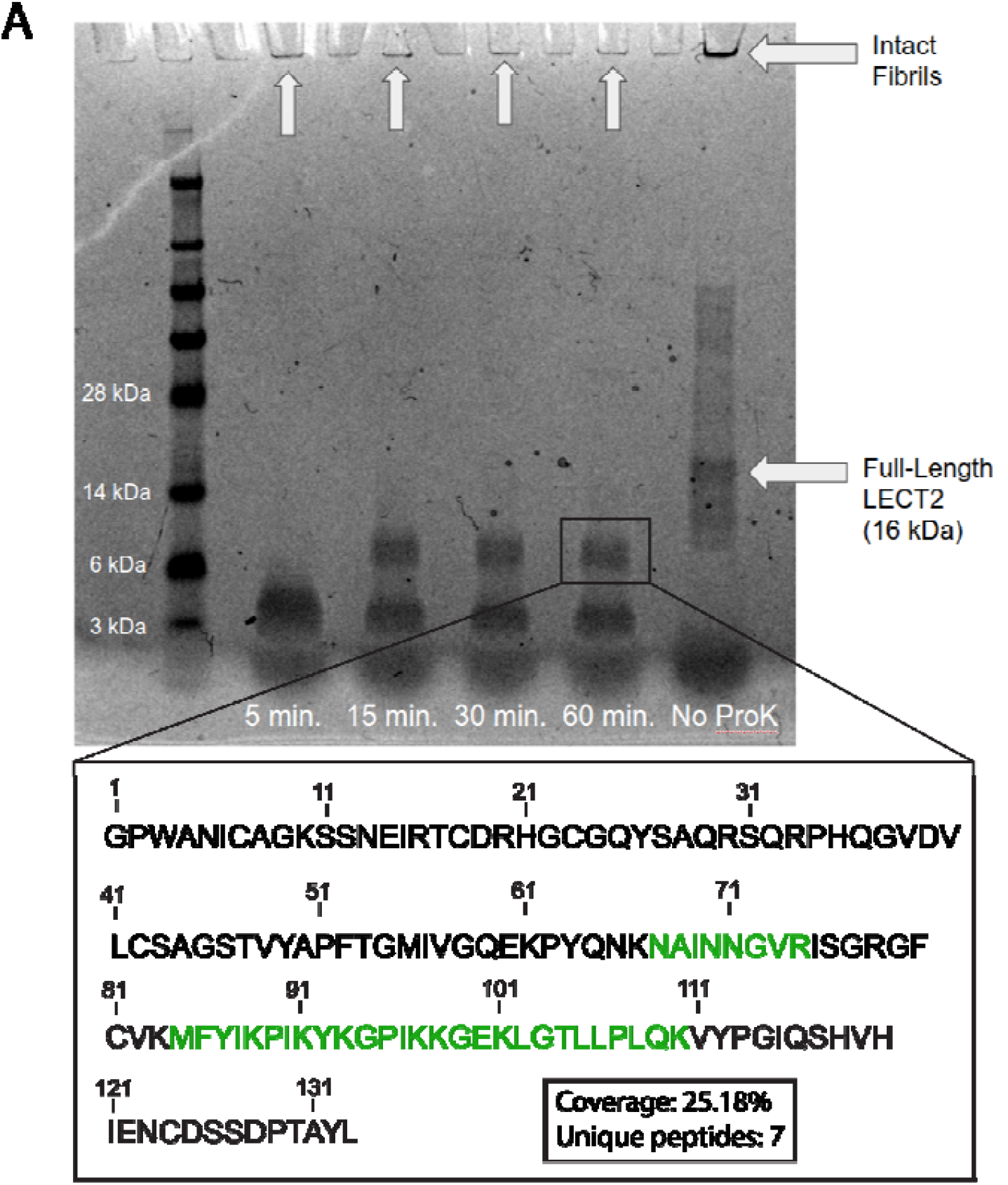
Mass spectrometry of Proteinase K digested fibrils. A) SDS-PAGE shows stable protein species present over one hour of Proteinase K digestion. Boxed band was excised and peptides were extracted for GeLC-MS/MS (inset) with sequence segments detected colored in green.

**Supplementary Table 2:**
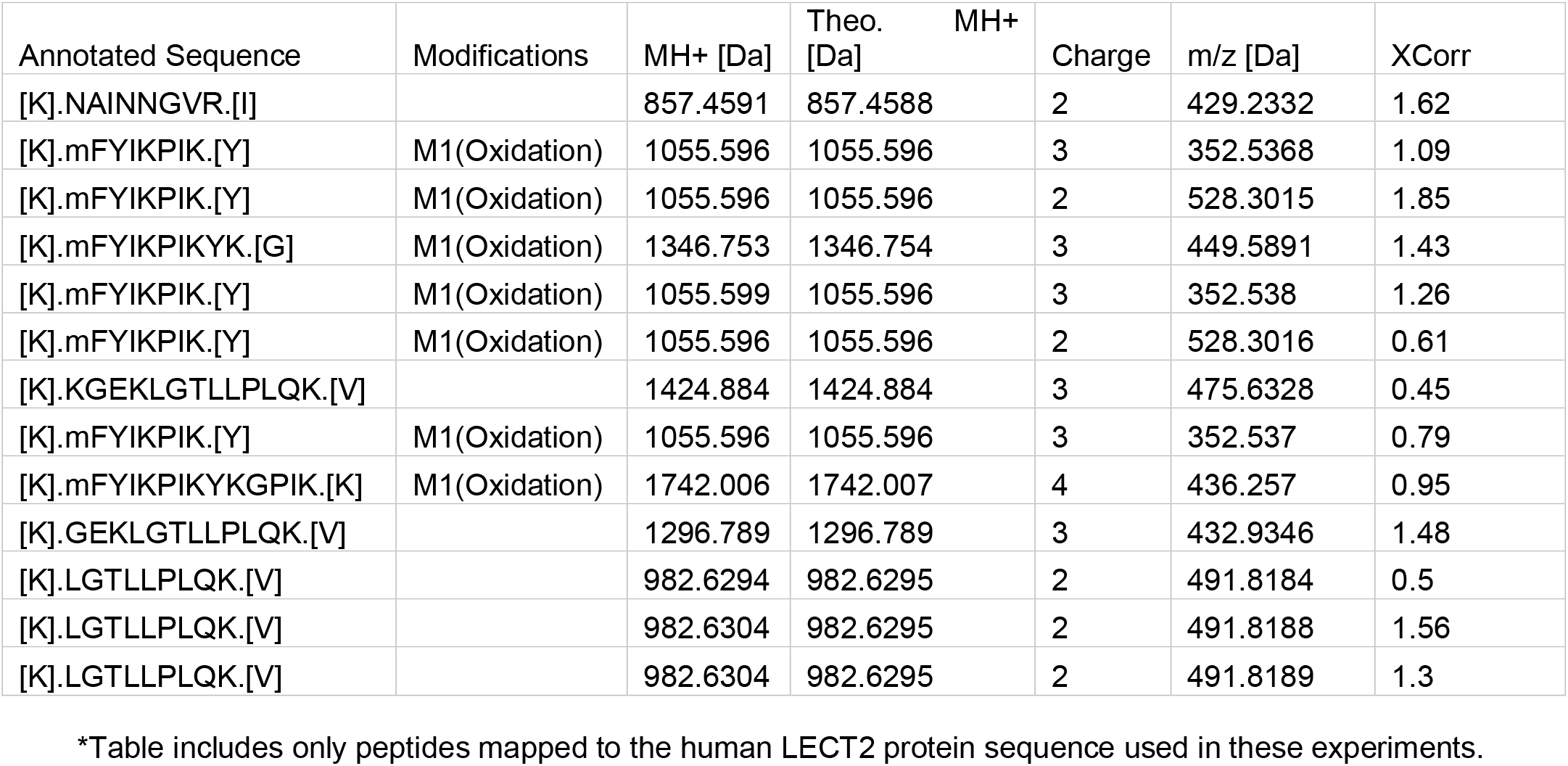
Summary of Proteinase K digested LECT2 peptides identified by GeLC-MS/MS.

## Supplementary Code

PyRosetta Sequence Threading

~~~
import os
import numpy as np
import pandas as pd
from pathlib import Path
import csv
from pyrosetta import *
from pyrosetta.rosetta import *
from pyrosetta.teaching import *
from pyrosetta.toolbox import *
from rosetta.protocols.cryst import *
from rosetta.protocols.rosetta_scripts import *
np.set_printoptions(threshold=sys.maxsize)
init(‘-ignore_unrecognized_res -load_PDB_components false -ignore_zero_occupancy false’)
#####################################################
### INPUTS ###
#####################################################
### protein name
protName = ‘lect2’
### to build symmetry pose
symmInfo = ‘lect2.symm’
pose_file = ‘lect2_INPUT.pdb’
### start/end residues in pose numbering for base layer of symmetric pose
base_start = 85
base_end = 105
### for job distribution
job = int(sys.argv[1])
### sliding window size
window_size = 21
### full sequence txt file to be threaded
seq_file = ‘seq.txt’
### output name for scores file
scores_output = ‘lect2_scores.csv’
#####################################################
### PREPROCESSING ###
#####################################################
### pre prosess sequence file
seq = Path(seq_file).read_text()
seq = seq.replace(‘\n’, ‘‘)
seq_list = list(seq)
#####################################################
### FUNCTIONS ###
#####################################################
scorefxn = get_fa_scorefxn()
def symmetrize_pose(pose):
 “““ set up symmetric pose “““
 pose_symm_data = core.conformation.symmetry.SymmData(pose.total_residue(), pose.num_jump())
 pose_symm_data.read_symmetry_data_from_file(symmInfo)
 core.pose.symmetry.make_symmetric_pose(pose, pose_symm_data)
 sym_info = pose.conformation().Symmetry_Info()
print(“AssymUnit? equivalent_res”)
for i in range(1, pose.size()+1):
 print(i, sym_info.bb_is_independent(i), sym_info.bb_follows(i))
print(“Total Subunits:”, sym_info.subunits())
return pose
def def_windows(x, wnd_sz):
 “““ define sequence positions for each window “““
 window_list = []
start_pos = 0
end_pos = start_pos + wnd_sz
num_windows = len(x) - wnd_sz + 1
for j in range(0, num_windows):
window_tmp = []
for i in range(start_pos, end_pos):
 window_tmp.append(x[i])
start_pos += 1
end_pos += 1
window_list.append(window_tmp)
return window_list
def thread(pose, x, base_start, base_end):
“““ perform threading
x = sequence window list “““
j = 0
for i in range(base_start, base_end+1):
 mutate_residue(pose, i, x[job][j])
 j += 1
def FRwDensity(pose, rpt_num):
 “““ set up and perform FastRelax with cryoEM density map “““
## set up density FastRelax
setup_dens = XmlObjects.static_get_mover(‘<LoadDensityMap name = “loaddens” mapfile=“postprocess.mrc”/>‘)
setup_dens.apply(pose)
setup_dens_pose = rosetta.protocols.electron_density.SetupForDensityScoringMover()
setup_dens_pose.apply(pose)
## set up score function with correct weights
score = get_score_function()
score_dens_cart = create_score_function(‘ref2015_cart’)
score_dens_cart.set_weight(rosetta.core.scoring.elec_dens_fast, 25)
score_dens_cart.set_weight(fa_elec, 1.5)
## set up fast relax parameters
mmf = pyrosetta.rosetta.core.select.movemap.MoveMapFactory()
mmf.all_bb(setting=True)
mmf.all_bondangles(setting=True)
mmf.all_chi(setting=True)
mmf.all_jumps(setting=True)
mmf.set_cartesian(setting=True)
FR = pyrosetta.rosetta.protocols.relax.FastRelax(scorefxn_in=score_dens_cart, standard_repeats=1)
FR.cartesian(True)
FR.set_movemap_factory(mmf)
FR.min_type(“lbfgs_armijo_nonmonotone”)
## apply FR, score relaxed pdb, and output file
FR.apply(pose)
output_name = ‘lect2_FR_thread_’ + str(job) + ‘_’ + str(rpt_num) + ‘.pdb’
pose.dump_pdb(‘./output/’ + output_name)
scores = output_name, score_dens_cart(pose), scorefxn(pose) return scores
#####################################################
### RUN ###
#####################################################
window_list = def_windows(seq_list, window_size) ## build list of sequence windows
seq_list_df = pd.DataFrame(window_list) ## convert to df
for i in range(0, 3):
pose = pose_from_pdb(pose_file) ## create pose object
pose = symmetrize_pose(pose) ## symmetrize the pose
thread(pose, window_list, base_start, base_end) ## perform threading; mutate side chains on
pose object to reflect the current sequence window
scores = FRwDensity(pose, i) ## perform FastRelax on current sequence window
~~~

